# Early Proteomic and Metabolic Signatures of Liver and Eye in OAT-Deficient Mice

**DOI:** 10.1101/2025.09.10.675383

**Authors:** Artjola Puja, Rong Xu, Isabella Mascari, Tuan Ngo, Ying Zhang, Qingyan Wang, Meghashri Saravanan, Jianhai Du

## Abstract

Ornithine aminotransferase (OAT) links the urea cycle, TCA cycle, and amino acid metabolism by interconverting ornithine to pyrroline-5-carboxylate and glutamate. Mutations in *OAT* cause hyperornithinemia and predominantly affect the eye, leading to gyrate atrophy of the choroid and retina (GA), a rare inherited blinding disorder. To understand the early molecular changes that make the eye susceptible to damage, we performed quantitative proteomic and metabolomic profiling of liver, retina, and retinal pigment epithelium and choroid (RPE/Cho) from OAT-deficient (*Oat^rhg^*) mice prior to detectable vision impairment. In addition to reduced OAT expression and elevated ornithine, methylation-related metabolites such as N(6)-methyl-lysine were altered in all examined tissues of *Oat^rhg^*mice. In the liver, ornithine disposal through the urea cycle was enhanced, together with altered expression of detoxification enzymes and histone H2B proteins. In contrast, the retina had minimal proteomic changes but pronounced alterations in amino acid pathways supporting glutamate homeostasis. The RPE/Cho demonstrated the most extensive proteomic changes, particularly in mitochondrial metabolism, cytoskeleton, and extracellular matrix, along with reductions in metabolites involved energy metabolism and antioxidant capacity. Together, these findings highlight common and tissue-specific impacts of OAT on the liver and ocular tissues and provide insight into early molecular changes that contribute to the selective vulnerability of the eye in GA. Proteomics data are available via ProteomeXchange (PXD063614) and metabolomics data via MassIVE repository (MSV000101103).

## 1. Introduction

Ornithine aminotransferase (OAT) is a pyridoxal-5-phosphate (PLP)-dependent mitochondrial enzyme that catalyzes the reversible transamination of ornithine, a urea cycle intermediate, and α-ketoglutarate (α-KG) to generate pyrroline-5-carboxylate (P5C) and glutamate. P5C is a precursor for proline synthesis, while glutamate and α-KG feed into the tricarboxylic acid (TCA) cycle, positioning OAT at the intersection of nitrogen and energy metabolism (**Figure 1A**). Mutations in the *OAT* gene cause gyrate atrophy of the choroid and retina (GA), a rare autosomal recessive blinding disorder characterized by markedly elevated plasma ornithine levels [1–3]. The clinical presentation of GA is dominated by ocular pathology, with rare and mild neurological and muscular manifestations [4–6]. Ocular symptoms typically present in childhood with myopia and night blindness, followed by progressive chorioretinal atrophy and peripheral vision loss. Cataracts and macular edema may also occur, with variable rate of progression [7–10].

**Figure 1.**
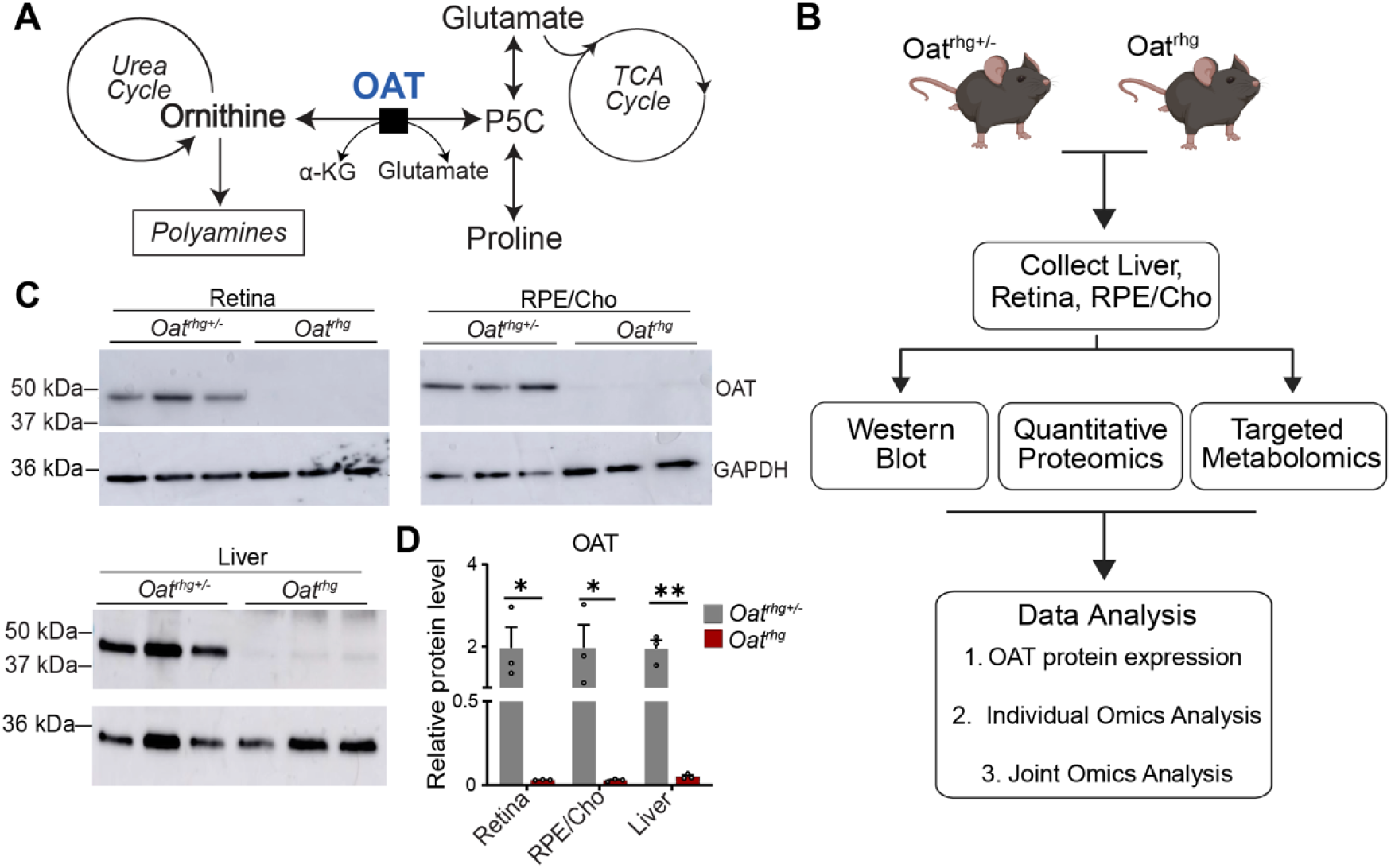
Experimental workflow and mouse model validation. **(A)** Illustration of the OAT reaction and connected metabolic pathways. **(B)** Schematic overview of the experimental workflow. **(C)** Western blot confirming deficiency of OAT protein expression in retina, RPE/Cho, and liver tissues in *Oat^rhg^* compared to *Oat^rhg+/-^*and **(D)** quantification normalized to GAPDH loading control (N=3). p < 0.05. RPE/Choroid (RPE/Cho); α-ketoglutarate (α-KG); Pyrroline-5-carboxylate (P5C); Ornithine aminotransferase (OAT); Glyceraldehyde 3-phosphate dehydrogenase (GAPDH), Tricarboxylic acid cycle (TCA cycle).

A distinctive biochemical feature of GA in adult patients is a 10- to 20-fold elevation of plasma ornithine. High ornithine has long been considered toxic to the eye; however, the underlying mechanisms remain poorly understood [11–14]. Importantly, plasma ornithine levels do not consistently correlate with disease severity, and hyperornithinemia alone, as observed in hyperornithinemia-hyperammonemia-homocitrullinuria (HHH) syndrome, does not affect vision [15,16]. In addition to ornithine, levels of multiple plasma amino acids are altered in GA patients, including lysine, creatine, proline, and histidine [7,12,17]. Plasma lysine is consistently decreased, likely due to the competition with ornithine and arginine for shared renal transporters, as lysinuria has been reported in some patients [12,18,19]. Moreover, plasma creatine is reduced, reflecting the inhibition of arginine:glycine amidinotransferase (AGAT) by accumulated ornithine [20]. This reduction may contribute to extraocular manifestations in GA, particularly affecting muscles and the brain [20–23]. In addition, ornithine can be funneled through ornithine decarboxylase (ODC) into polyamine synthesis, a process that consumes S-adenosylmethionine (SAM), a universal methyl donor for the methylation of DNA, histones, and small molecules [24,25]. SAM links methylation to both redox and nucleotide metabolism by generating homocysteine for glutathione synthesis and interacting with the folate cycle that supports purine production. Elevated polyamines have been shown to be toxic to retinal pigment epithelium (RPE) cells, induce RPE dysfunction [26,27], and increased polyamine levels have been reported in plasma and urine of some GA patients.

Currently, there is no treatment for GA. Disease management aims to reduce systemic ornithine with an arginine-restricted diet and support downstream pathways using supplements such as proline, creatine, lysine, and pyridoxine (vitamin B6) [23,28–33]. These interventions can slow disease progression; however, most patients ultimately become blind. Recent work has shown that liver-targeted gene therapy reduces systemic ornithine and partially improves retinal outcomes in OAT-deficient mice [34]. Importantly, combined restoration of OAT in both liver and retina provides greater improvements in retinal structure and function than targeting either tissue alone [35], demonstrating that the loss of OAT activity in both tissues contributes to the disease phenotype. However, how OAT deficiency and hyperornithinemia affect these tissues at the molecular level, particularly in driving vision impairment, remains undefined.

To identify early molecular changes associated with OAT deficiency prior to overt retinal degeneration, we performed integrated proteomic and metabolomic profiling of liver, retina, and RPE/choroid (RPE/Cho) from young adult *Oat^rhg^* mice (*Oat*^G353A/G353A^), which recapitulate the key biochemical and ocular features of GA [36]. We found that OAT deficiency triggers tissue-specific responses, affecting amino acid metabolism, the urea cycle, SAM-dependent pathways, detoxification, and remodeling of histone and extracellular matrix (ECM) proteins. These findings provide a mechanistic framework of how OAT deficiency disrupts cellular homeostasis in the liver and eye.

## 2. Materials and methods

### 2.1. Animals

All animal procedures were conducted in accordance with National Institutes of Health guidelines and were approved by the Institutional Animal Care and Use Committee (IACUC) of West Virginia University. All animals were housed under standard conditions with *ad libitum* access to standard chow and water. *Oat* homozygous mutants (*Oat^rhg^)* mice were obtained from the Jackson Laboratory and bred onto a C57BL/6J background for more than three generations before use in experiments. Heterozygous *Oat^rhg+/-^* mice are used as controls in this study and retain OAT activity comparable to wild-type controls [36]. Adults at 3.5 months of age from both genotypes were used for experiments. Based on prior reports, retinal and RPE/Cho structure and function are normal at these ages, allowing us to analyze molecular changes preceding visual dysfunction [34,36]. Genotyping was performed by Transnetyx, Inc. (Cordova, TN; see **Table S1**).

### 2.2. Tissue collection

Samples for metabolomics and proteomics were collected around noon from separate sets of adult *Oat^rhg^* and littermate *Oat^rhg+/-^* control mice of both sexes. Prior to tissue collection, all animals were transferred to a procedure room and acclimated for one hour. Mice were euthanized via cervical dislocation. Their eyes were immediately enucleated, and the surrounding muscle and fat were carefully removed. The neural retina and RPE/Cho were then isolated in cold HBSS by dissecting the anterior segments as previously described [37]. Because the RPE is difficult to separate from its tightly bound choroid, and the choroid plays an important role in RPE function, we analyzed the RPE/choroid complex (RPE/Cho) as in our previous studies [38]. A small section from the right lobe of the liver was excised and rapidly rinsed in cold PBS. All collected tissues were flash-frozen in liquid nitrogen.

### 2.3. Proteomics sample preparation

Proteins from the retina, RPE/Cho, and liver were extracted using radioimmunoprecipitation assay (RIPA) buffer with protease and phosphatase inhibitors (5 mg/mL). The tissue lysates were centrifuged at 12,000 rpm for 15 minutes at 4°C, and the supernatant was used to measure protein concentration by BCA assay. The quantitative proteomic analysis was performed by IDeA National Resource for Quantitative Proteomics (Little Rock, Arkansas).

### 2.4. Quantitative proteomics

Total protein from each sample was reduced, alkylated, and purified by chloroform/methanol extraction prior to digestion with sequencing-grade modified porcine trypsin (Promega). Tryptic peptides were then separated by reverse phase XSelect CSH C18 2.5 um resin (Waters) on an in-line 150 x 0.075 mm column using an UltiMate 3000 RSLCnano system (Thermo). Peptides were eluted using a 60 min gradient from 98:2 to 65:35 buffer A:B ratio (Buffer A contains 0.1% formic acid, 0.5% acetonitrile; Buffer B contains 0.1% formic acid, 99.9% acetonitrile). Eluted peptides were ionized by electrospray (2.2 kV) followed by mass spectrometric analysis on an Orbitrap Exploris 480 mass spectrometer (Thermo). To assemble a chromatogram library, six gas-phase fractions were acquired on the Orbitrap Exploris with 4 m/z DIA spectra (4 m/z precursor isolation windows at 30,000 resolution, normalized AGC target 100%, maximum inject time 66 ms) using a staggered window pattern from narrow mass ranges using optimized window placements. Precursor spectra were acquired after each DIA duty cycle, spanning the m/z range of the gas-phase fraction (i.e. 496-602 m/z, 60,000 resolution, normalized AGC target 100%, maximum injection time 50 ms). For wide-window acquisitions, the Orbitrap Exploris was configured to acquire a precursor scan (385-1015 m/z, 60,000 resolution, normalized AGC target 100%, maximum injection time 50 ms) followed by 50x 12 m/z DIA spectra (12 m/z precursor isolation windows at 15,000 resolution, normalized AGC target 100%, maximum injection time 33 ms) using a staggered window pattern with optimized window placements. Precursor spectra were acquired after each DIA duty cycle.

### 2.5. Proteomics data analysis

Following data acquisition, data were searched using an empirically corrected library against the UniProt Mus musculus database (Proteome ID: UP000000589, 2nd version of 2023), and a quantitative analysis was performed to obtain a comprehensive proteomic profile. Proteins were identified and quantified using EncyclopeDIA [39] and visualized with Scaffold DIA using 1% false discovery thresholds at both the protein and peptide level. Protein MS2 exclusive intensity values were assessed for quality using ProteiNorm [40]. The data were normalized using Cyclic Loess [41] to correct for systemic variation. Differential expression was assessed using proteoDA, which implements linear models (limma) with empirical Bayes (eBayes) with empirical Bayes (eBayes) smoothing to the standard errors [42,43]. Proteins with an adjusted p-value < 0.05 and a fold change (FC) > 2 were considered significant.

### 2.6. Proteomic pathway analysis

Pathway enrichment and functional annotation of differentially expressed proteins were performed using STRING (v12.0) [44] and DAVID [45]. Statistical significance was defined as |log2fold change| ≥ 1 and a p-value < 0.05. Venn diagrams were generated using InteractiVenn [46] to visualize protein overlaps across tissues. Metabolic proteins were identified by intersecting significantly altered proteins with a previously published metabolic gene set [47], followed by manual curation using UniProt and NCBI/PubMed. For each protein, literature validation was performed using Boolean query strings combining the protein identifier with keywords for ocular tissues, liver, and metabolic function (e.g., “[protein name] AND (retina OR RPE OR liver OR metabolism)”).

### 2.7. Metabolite analysis and data processing

Metabolites were extracted from tissues by homogenization in ice-cold 80% (v/v) methanol and from plasma in ice-cold 100% methanol, as previously described [48]. Homogenates were centrifuged at 13,000 rpm for 15 min at 4°C, and supernatants were vacuum-dried (Eppendorf Vacufuge Plus) and analyzed in liquid chromatography-mass spectrometry (LC–MS). Targeted metabolite profiling was performed on a Shimadzu Nexera UHPLC system coupled to an AB Sciex QTRAP 5500 mass spectrometer (AB Sciex, Toronto, ON, Canada) using a multiple reaction monitoring (MRM) method as previously described [47,49]. Metabolite peaks were identified and quantified using MultiQuant Software (v3.0). Statistical analysis, volcano plots (thresholds: P < 0.05 and fold change > 1.3), Partial Least Squares Discriminant Analysis (PLS-DA), and pathway enrichment analysis were performed using MetaboAnalyst 6.0 (https://www.metaboanalyst.ca). Raw mass spectrometry data have been deposited to MassIVE repository (MSV000101103).

### 2.8. Western Blot

Immunoblots were performed as previously described [50]. Briefly, tissue samples containing 20Lμg protein were separated in Mini-PROTEAN TGX precast gels, transferred onto 0.22 μm nitrocellulose membranes, and blocked with 5% non-fat milk in 1× Tris-Buffered Saline with 0.1% Tween-20 (TBST). Membranes were incubated overnight at 4°C with primary antibodies (1:1000 dilution; see **Table S1**) and then in horseradish peroxidase (HRP)-conjugated secondary antibodies (1:2000) for 1 hour, followed by visualization by chemiluminescence.

### 2.9. Statistics

Ornithine ion abundances in *Oat^rhg^* mice were divided by those from *Oat^rhg+/-^* mice to calculate fold changes (FC). Data were graphed using GraphPad Prism (v9.5.1). Statistical significance was determined using multiple unpaired t-tests with Bonferroni correction method, with pL<L0.05 considered significant. All data are presented as the mean ± SD.

## 3. Results

### 3.1. OAT deficiency causes ocular and liver ornithine accumulation before visual decline in Oat^rhg^ mice

To capture early molecular changes that precede overt degeneration, we performed quantitative proteomics and targeted metabolomics on the retina, RPE/Cho, and liver from 3.511month11old *Oat^rhg^* mice and *Oat^rhg+/-^* littermate controls, tissues with normally high OAT activity [34,51] (**Figure 1B**). At this age, *Oat^rhg^* mice appear healthy, with body size comparable to *Oat^rhg+/-^*controls. Although loss of OAT enzymatic activity in *Oat^rhg^* mice has been established [36], OAT protein expression in ocular tissues has not been assessed. Western blot analysis demonstrated a complete absence of OAT protein in retina, RPE/Cho, and liver, confirming its loss in mutant mice (**Figure 1C, D, S1**). Consistent with previous reports, plasma ornithine levels were more than 6-fold higher in adult *Oat^rhg^* mice [52–54]. Ornithine levels in liver, retina, and RPE/Cho were also dramatically increased, exceeding those in controls by over 20-fold (**Figure S2**). Despite systemic and tissue ornithine accumulation, electroretinogram (ERG) recordings showed normal visual function in 3.511month11old *Oat^rhg^* mice, indicating that retinal degeneration had not yet developed at this stage (**Figure S3**). These data confirm global ornithine accumulation and lack of OAT protein in ocular and liver tissues, establishing this pre-degenerative stage as suitable for multi11omics analyses to define early molecular changes that may underlie the later selective vulnerability of the eye.

### 3.2. OAT deficiency alters liver proteome in multiple pathways beyond ornithine metabolism

To determine the molecular changes underlying OAT deficiency in the liver, we conducted quantitative proteomic profiling of tissues from adult *Oat^rhg^*and *Oat^rhg+/-^* mice. The proteomics analysis identified 62 differentially expressed (DE) proteins in adult *Oat^rhg^* mouse liver compared to controls (**Figure 2A**). As expected, OAT was the most significantly changed protein with an approximately 9-fold decrease. Functional categorization of these DE proteins revealed a significant enrichment in metabolism (34%), followed by transcription (18%), cytoskeleton (13%), immunity (11%), and signal transduction (10%) (**Figure 2B**). Subcellular localization analysis showed that these proteins were predominantly associated with the cytoplasm, endoplasmic reticulum (ER), nucleus, and membranes, while mitochondrial and lysosomal proteins were less affected (**Figure 2C**). Gene ontology (GO) analysis revealed that the top five enriched biological pathways in the OAT-deficient liver were nucleosome assembly, steroid, alcohol, and xenobiotic metabolism, as well as oxidative demethylation (**Figure 2D**).

**Figure 2.**
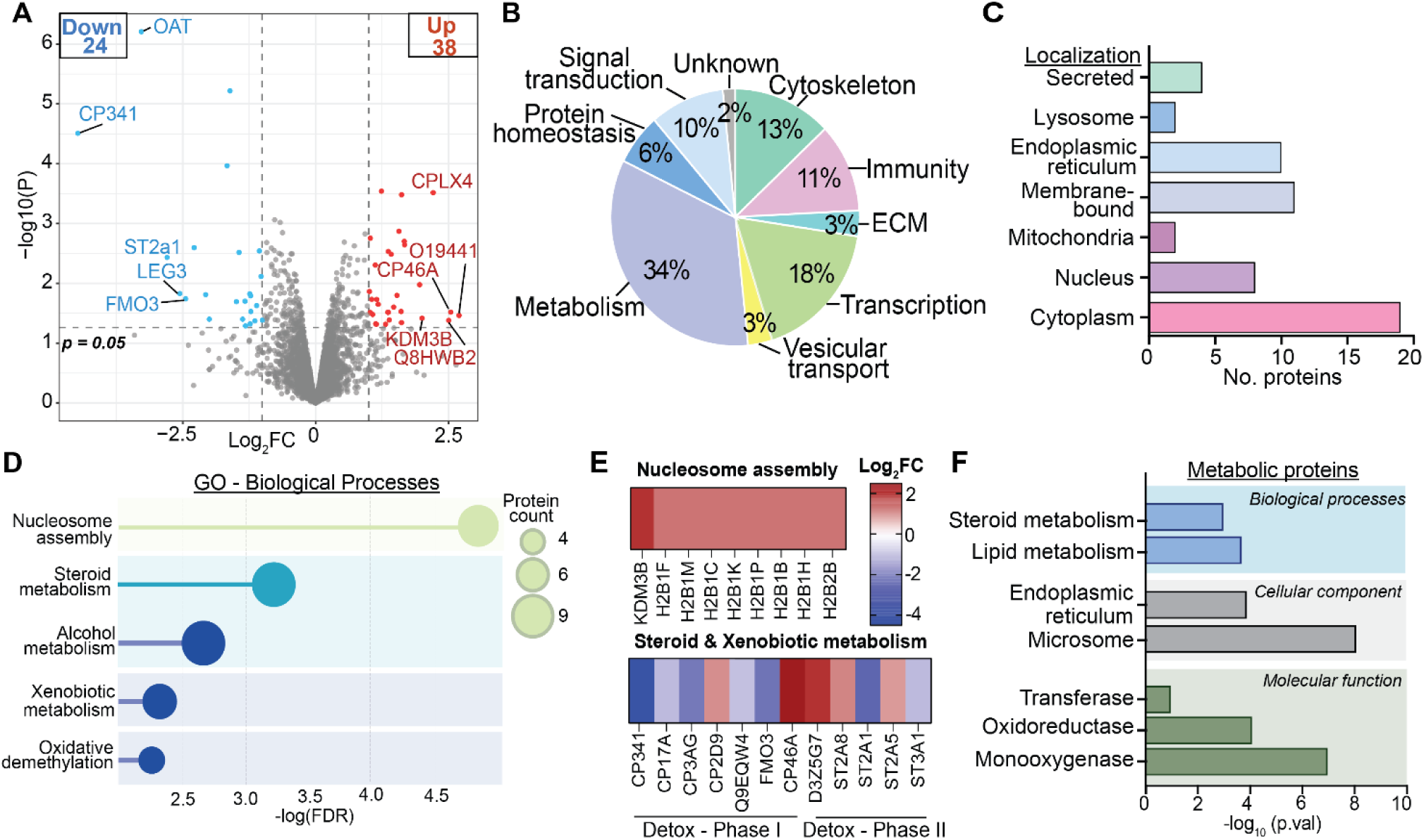
Analysis of DE proteins in *Oat^rhg^* liver tissues compared to controls. **(A)** Volcano plot showing top 5 up and downregulated proteins in *Oat^rhg^* liver and **(B)** their biological function classification based on manual search in UniProt and NCBI/PubMed. **(C)** Subcellular localization of DE proteins in the liver based on UniProt annotations. **(D)** Top five enriched biological processes identified from the analysis of DE proteins using STRING. **(E)** Extended view of enriched pathways and their associated proteins. **(F)** Functional annotation analysis of enriched metabolic proteins in the liver. Detox-Phase I (oxidation/reduction by cytochrome P450 enzymes); Detox-Phase II (conjugation reactions such as sulfation and ester hydrolysis); Major Histocompatibility Complex (MHC). N=3. |log_2_fold change| ≥ 1, p-value < 0.05.

To further define these changes, we examined key altered proteins within these categories (**Figure 2E**). We found that several histone H2B variants (H2B1F, H2B1M, H2B1C) and demethylases (KDM3B) were upregulated, suggesting chromatin remodeling and transcriptional alterations. In addition, we observed substantial disruption of the Phase I and II detoxification pathways, involving cytochrome P450 enzymes (CP341, CP17A, CP3AG, CP341) and sulfotransferases (ST2A1, ST2A5, ST2A8, ST3A1), which are critical for the metabolism of drugs, steroids, bile acids, and fatty acids. Flavin-containing monooxygenase 3 (FMO3), a key hepatic monooxygenase, was also markedly reduced, potentially altering the detoxification capacity of the liver in *Oat^rhg^* mice. To refine the metabolic changes in the liver, we next performed GO analysis specifically on DE metabolic proteins, examining biological processes, cellular components, and molecular functions (**Figure 2F**). The analysis showed that altered metabolic proteins in *Oat^rhg^* liver were enriched in steroid and lipid metabolism, mainly localized to the ER and microsomes, and associated with oxidoreductase, monooxygenase, and transferase activities. Taken together, these results illustrate that OAT deficiency in the adult liver extends beyond amino acid transamination, potentially impacting diverse pathways including steroid and lipid metabolism, detoxification, and transcriptional regulation.

### 3.3. Metabolomic analysis reveals early alterations of metabolic processes in Oat^rhg^ liver

Targeted metabolomics quantified 144 metabolites in liver from *Oat^rhg^*and *Oat^rhg+/-^* mice. The Partial Least Squares Discriminant Analysis (PLS-DA) showed clear separation between genotypes, indicating a distinct metabolic profile in *Oat^rhg^* liver (**Figure 3A**). Statistical comparison between groups identified 7 significantly altered metabolites, 3 increased and 4 decreased in *Oat^rhg^* compared to *Oat^rhg+/-^*, as shown in **Figure 3B**. Consistent with the canonical role of OAT in ornithine catabolism, targeted metabolomics revealed marked accumulation of ornithine and citrulline in *Oat^rhg^* liver, together with increased betaine and decreased glutathione, lactate, fructose, and N(6)-methyl-lysine. KEGG enrichment analysis highlighted the involvement of these metabolites in arginine and proline biosynthesis, one-carbon metabolism, glutathione metabolism, fructose and mannose metabolism, glycine/serine/threonine metabolism, and fatty acid degradation (**Figure 3C**). Together, these results reveal that loss of OAT activity triggers a broad metabolic reprogramming in the liver, in which excess ornithine is redirected into the urea cycle and polyamine synthesis, with secondary consequences for energy metabolism, S-adenosyl methionine (SAM)-dependent methylation, and glutathione-mediated redox homeostasis (**Figure 3D**).

**Figure 3.**
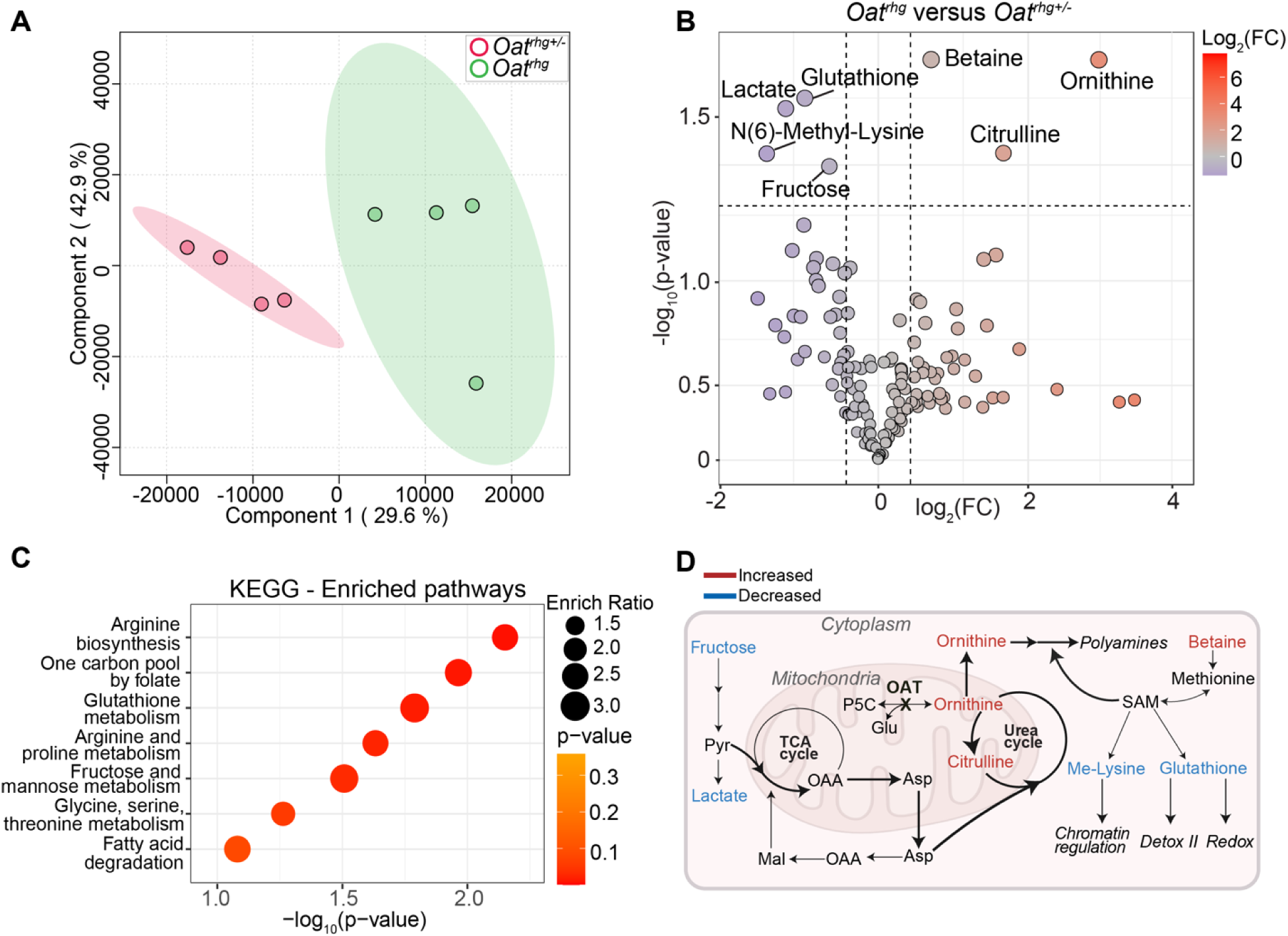
Metabolite analysis in liver tissues of *Oat^rhg^* compared to control mice. **(A)** Partial least squares discriminant analysis (PLS-DA) of liver metabolomics distinguishes *Oat^rhg^*from *Oat^rhg+/-^* mice. **(B)** Volcano plot of liver metabolomics showing increased (right) and decreased (left) metabolites in *Oat^rhg^* relative to *Oat^rhg+/-^*. **(C)** KEGG pathway enrichment analysis of liver metabolomics comparing *Oat^rhg^* and *Oat^rhg+/^*^-^. **(D)** Proposed model of metabolic alterations in *Oat^rhg^* liver. In the liver, excess ornithine is redirected into the urea cycle, increasing the demand for cytosolic aspartate and competing with the malate-aspartate shuttle. Fructose-derived pyruvate provides an alternative carbon source to sustain both TCA cycle activity and shuttle function. Excess ornithine may also be utilized for polyamine synthesis, affecting the SAM availability and downstream pathways. Red indicates increased, blue decreased in *Oat^rhg^*relative to controls. Darker arrows indicate putative enhanced metabolic flux. Pyruvate (Pyr); Aspartate (Asp); Oxaloacetate (OAA); Malate (Mal); Pyrroline-5-carboxylate (P5C); Ornithine aminotransferase (OAT); S-adenosyl methionine (SAM); Me-Lysine (N6-Methyl-Lysine). N=4. Fold change ≥1.3, p-value < 0.05.

### 3.4. OAT deficiency induces early disruption of retinal amino acid metabolism

Proteomic analysis of retina identified only 5 significantly DE proteins in *Oat^rhg^* mice compared to controls at this early stage. Among these, 3 were downregulated, including OAT, and 2 upregulated (**Figure 4A**). These proteins were primarily associated with cytoskeletal regulation (LIMA1, ROCK1) and synapse modulation (CAD13).

**Figure 4.**
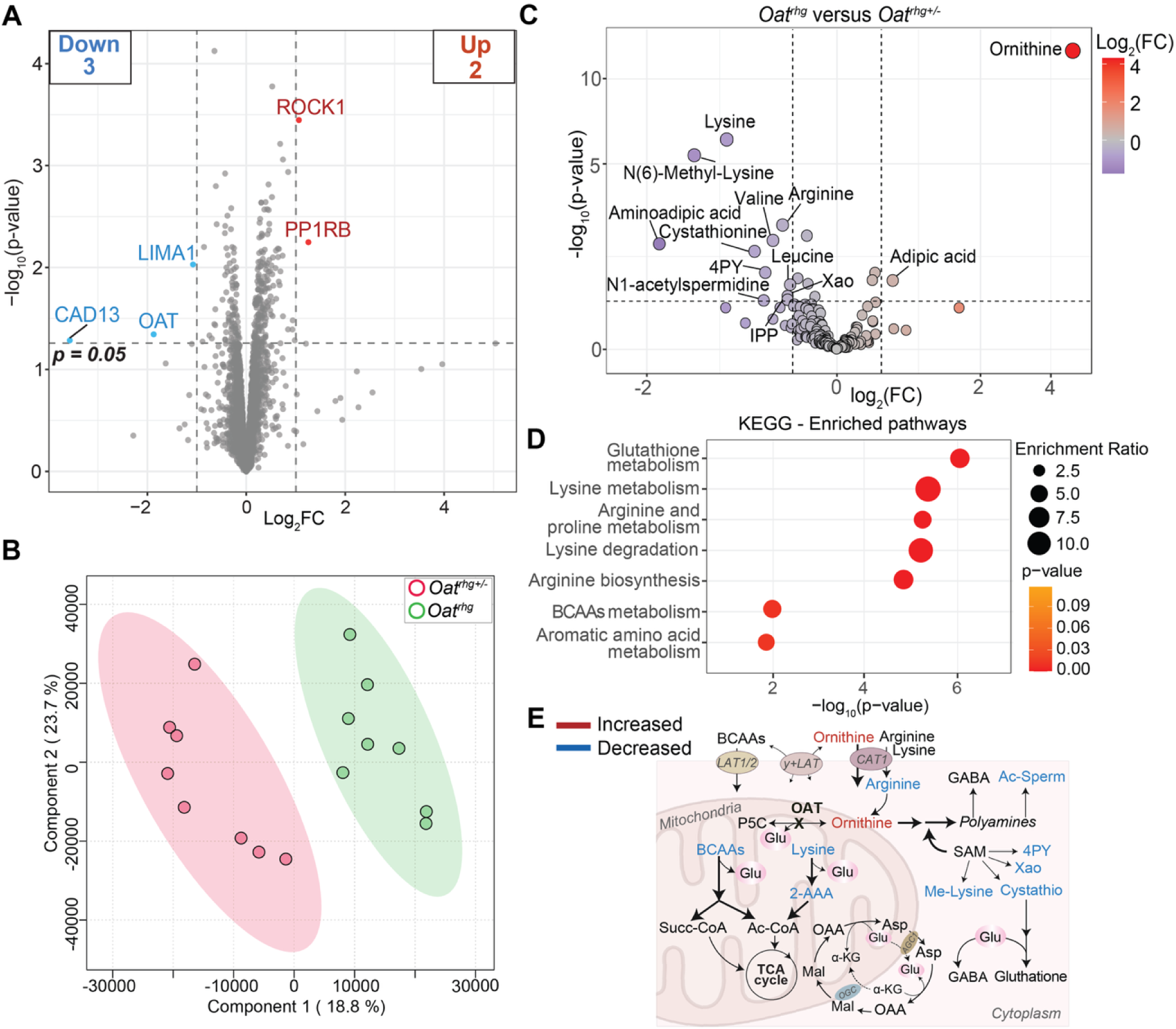
Proteome and metabolome changes in *Oat^rhg^* mouse retinas. **(A)** Volcano plot showing top 5 DE proteins in *Oat^rhg^* retina (N=3, |log_2_fold change| ≥ 1, p-value < 0.05). **(B)** Partial least squares discriminant analysis (PLS-DA) of retina metabolomics distinguishes *Oat^rhg^*from *Oat^rhg+/-^* mice. **(C)** Volcano plot showing significantly increased (right) and decreased (left) metabolites in *Oat^rhg^* retina compared to *Oat^rhg+/-^*(N=4, fold change ≥1.3, p-value < 0.05). **(D)** KEGG pathway enrichment analysis of retinal metabolomics comparing *Oat^rhg^* and *Oat^rhg+/^*^-^**. (E)** Proposed model of metabolic alterations in *Oat^rhg^* retina. Loss of OAT activity removes a direct source for glutamate in the retina. To maintain glutamate homeostasis, the retina compensates by increasing degradation of lysine and BCAAs, to generate glutamate and TCA cycle intermediates while preserving the α-KG/glutamate pool required for the malate-aspartate shuttle. Elevated ornithine may compete with other cationic amino acids for shared transporters, affecting their cellular uptake. Red indicates increased and blue decreased metabolites in *Oat^rhg^* relative to controls. Darker arrows indicate putative enhanced metabolic flux or transport. Branched chain amino acids (BCAAs); Acetyl CoA (Ac-CoA); Glutamate (Glu); Succinyl-CoA (Succ-CoA); Pyrroline-5-carboxylate (P5C); Ornithine aminotransferase (OAT); 2-aminoadipic acid (2-AAA); S-adenosyl methionine (SAM); Gamma amino-butyric acid (GABA); N1-Acetylspermidine (Ac-Sperm); N1-Methyl-4-pyridone-3-carboxamide (4PY); Cystathionine (Cystathio); Xanthosine (Xao); N(6)-Methyl-Lysine (Me-Lysine); Oxaloacetate (OAA); Malate (Mal); α-ketoglutarate (α-KG); Aspartate (Asp); Gamma-aminobutyric acid (GABA); L-type amino acid transporter 1/2 (LAT1/2); Cationic amino acid transporter 1 (CAT1); Aspartate/glutamate carrier (AGC1); Oxoglutarate carrier (OGC). N=4. Fold change ≥1.3, p-value < 0.05.

PLS-DA confirmed clear separation between *Oat^rhg^* and controls in retinal metabolome (**Figure 4B**). Targeted metabolomics detected 126 metabolites in the retina, of which 11 were significantly altered in *Oat^rhg^*compared to *Oat^rhg+/-^* mice (2 increased and 9 decreased) (**Figure 4C**). Ornithine was increased by more than 19-fold, together with a modest increase in adipic acid, an intermediate of dicarboxylic fatty acid metabolism. In contrast, multiple amino acids, including lysine, arginine, valine, and leucine, were reduced, together with lysine-derived aminoadipic acid. Catabolism of these amino acids supports mitochondrial metabolism and provides glutamate, a key neurotransmitter and nitrogen donor in the retina [47,55–58]. Several metabolites linked to one-carbon and polyamine pathways were also decreased, including N(6)-methyl-lysine, N1-methyl-4-pyridone-3-carboxamide (4PY), cystathionine, xanthosine, and N1-acetylspermidine, indicating reduced methyl donor availability and altered nucleotide and redox metabolism. KEGG analysis showed enrichment in glutathione metabolism and multiple amino acid pathways, including arginine and proline, lysine, branched-chain (BCAAs), and aromatic amino acid metabolism (**Figure 4D**). Overall, despite minimal proteomic changes, early retinal metabolomics indicate that OAT deficiency reprograms retinal amino acid metabolism to sustain glutamate neurotransmitter pool and replenish TCA cycle intermediates (**Figure 4E**).

### 3.5. RPE/Cho proteomics reveal early mitochondrial and structural changes in OAT deficiency

Among the three analyzed tissues, the RPE/Cho showed the most pronounced early proteomic changes, with 254 DE proteins in *Oat^rhg^* compared with *Oat^rhg+/-^* mice (155 downregulated and 99 upregulated; **Figure 5A**). These alterations involved metabolism (38%), cytoskeleton (15%), and signaling pathways (13%), and a substantial upregulation of crystallin proteins (CRYAA, CRBB2) (**Figure 5A, B**). GO analysis indicated reduced mRNA processing, ER-Golgi trafficking, and mitochondrial bioenergetics (i.e., electron transport chain). In contrast, pathways related to lipid and fatty acid metabolism, as well as cellular transport processes, were upregulated (**Figure 5C**). Most of these DE proteins were localized to the cytoplasm, mitochondria, and ER (**Figure 5D**). Strikingly, we found that the expression of several ribosomal subunit proteins (RL21, RL32, RL34, RS21) and the mitochondrial ribosomal protein RT35 were significantly decreased (**Figure 5E**). The RPE has a robust mitochondrial metabolism that supports energy production and nutrient synthesis for local use and export to the retina [57,59,60]. Several mitochondrial proteins, including mitochondrial transporters (TOM22, MPC1), components of the electron transport chain (CX7A1, COX2, NDUS8, NDUF4), amino acid (P5CR2, KCRS) and fatty acid metabolism enzymes (CPT1B, ABHDB, ACPM), were downregulated in RPE/Cho, suggesting an early disruption of mitochondrial function. In contrast, crystallin family proteins (CRYAA, CRBB1, CRBB2) were upregulated, potentially reflecting cellular stress response, consistent with previous reports [61,62].

**Figure 5.**
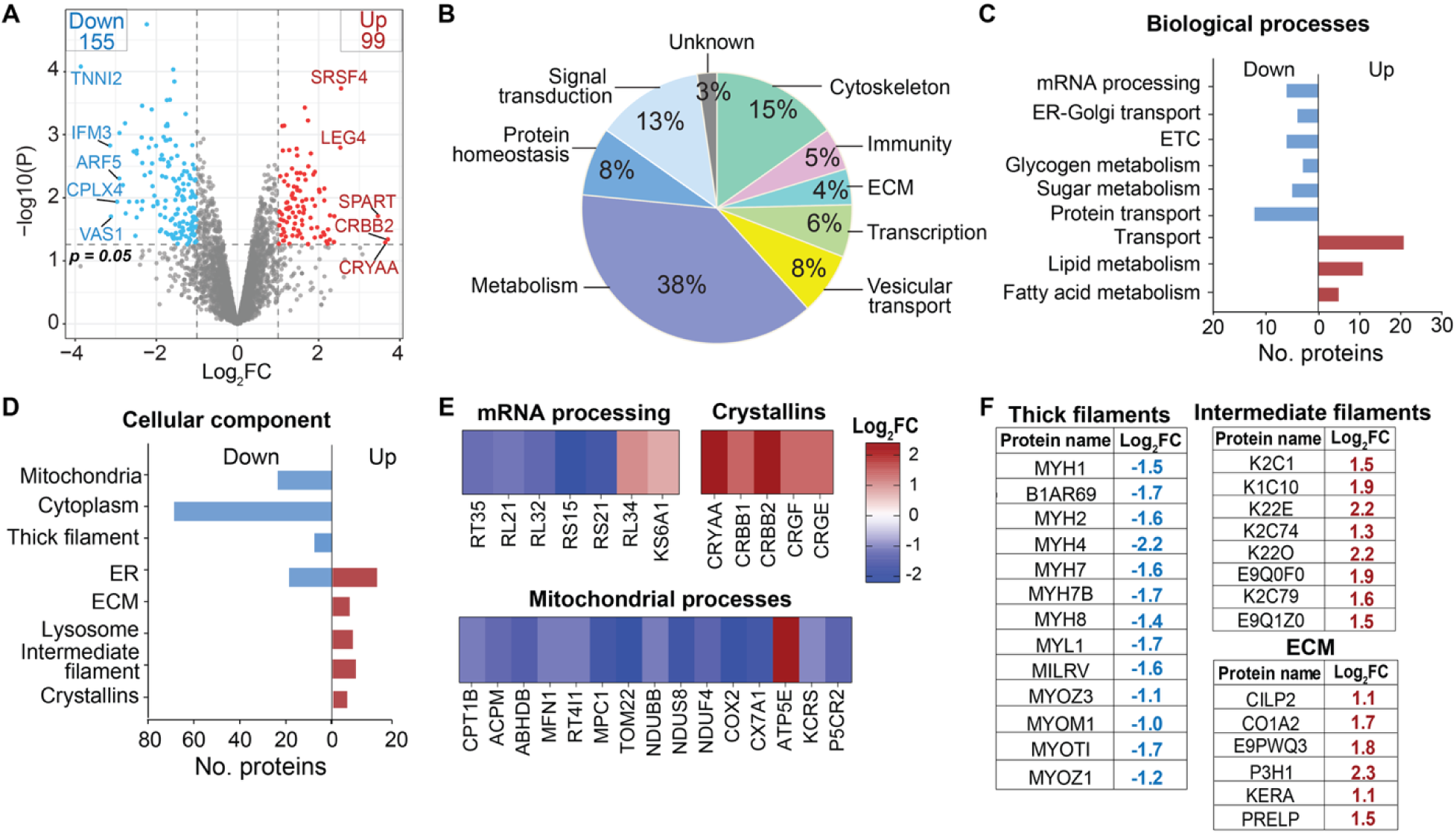
Proteomic changes in RPE/Cho in *Oat^rhg^* compared to controls. **(A)** Volcano plot highlighting top 5 up and downregulated proteins in *Oat^rhg^* RPE/Cho. **(B)** Pie chart shows enriched biological processes in RPE/Cho. **(C)** GO analysis of DE proteins in *Oat^rhg^* RPE/Cho showing enriched biological processes and **(D)** cellular components. Up and downregulated proteins were analyzed separately and are represented by red and blue bar graphs, respectively. **(E)** Heatmaps illustrate proteins associated with selected enriched biological and cellular processes. **(F)** List of DE proteins in *Oat^rhg^*associated with thick filaments (myosins), intermediate filaments (keratins), and extracellular matrix (ECM). Red and blue indicate upregulated and downregulated proteins, respectively. Endoplasmic reticulum (ER); Electron transport chain (ETC). N=3. |log_2_fold change| ≥ 1, p-value < 0.05.

Given the essential role of RPE metabolism in supporting retinal health [63,64], we further analyzed DE metabolic proteins in the RPE/Cho. GO enrichment analysis showed that these proteins were mainly involved in lipid and fatty acid metabolism, the respiratory chain, carbohydrate metabolism, and transport-related processes (**Figure S4**). The enriched proteins were predominantly localized to mitochondria, ER, microsomes, and inner mitochondrial membrane and were enriched for oxidoreductase, monooxygenase, hydrolase, and transferase activities. Interestingly, significant alterations were found in cytoskeletal, intermediate filament, and ECM proteins, with downregulation of myosins and upregulation of keratins and a few collagen proteins (**Figure 5F**). Overall, loss of OAT activity in RPE/Cho is associated with early proteomic changes characterized by impaired mitochondrial bioenergetics, reduced ribosomal proteins, and cytoskeletal–ECM remodeling.

### 3.6. OAT deficiency drives early metabolic changes in RPE/Cho linking amino acid metabolism to redox and structural processes

Targeted metabolomics quantified 136 metabolites in RPE/Cho from *Oat^rhg^*and *Oat^rhg+/-^* mice. PLS-DA showed that *Oat^rhg^*and control RPE/Cho formed distinct clusters, indicating that metabolite changes clearly separated the two groups (**Figure 6A**). Statistical analysis identified 7 significantly altered metabolites, of which 2 were increased and 5 decreased in *Oat^rhg^* RPE/Cho (**Figure 6B**). Ornithine was increased by more than 20⍰fold, accompanied by a more than 2⍰fold elevation in aminoadipic acid. Several metabolites linked to S-adenosyl methionine (SAM)-dependent pathways, including creatinine, 4PY, and N(6)⍰methyl⍰lysine, were reduced, consistent with the retina and liver, indicating limited methylation capacity. Decreased carnosine and myo-inositol further suggest disruption of antioxidant defense and phosphoinositide-related pathways, which are important for membrane dynamics and cellular structure [65–69]. KEGG enrichment analysis identified that OAT deficiency affects glutathione metabolism, arginine and proline metabolism, inositol phosphate pathway, and others, as shown in **Figure 6C**. Together, these results reveal that loss of OAT activity in the RPE/Cho perturbs amino acid metabolism, antioxidant response, and processes that support cellular structure and dynamics (**Figure 6D**).

**Figure 6.**
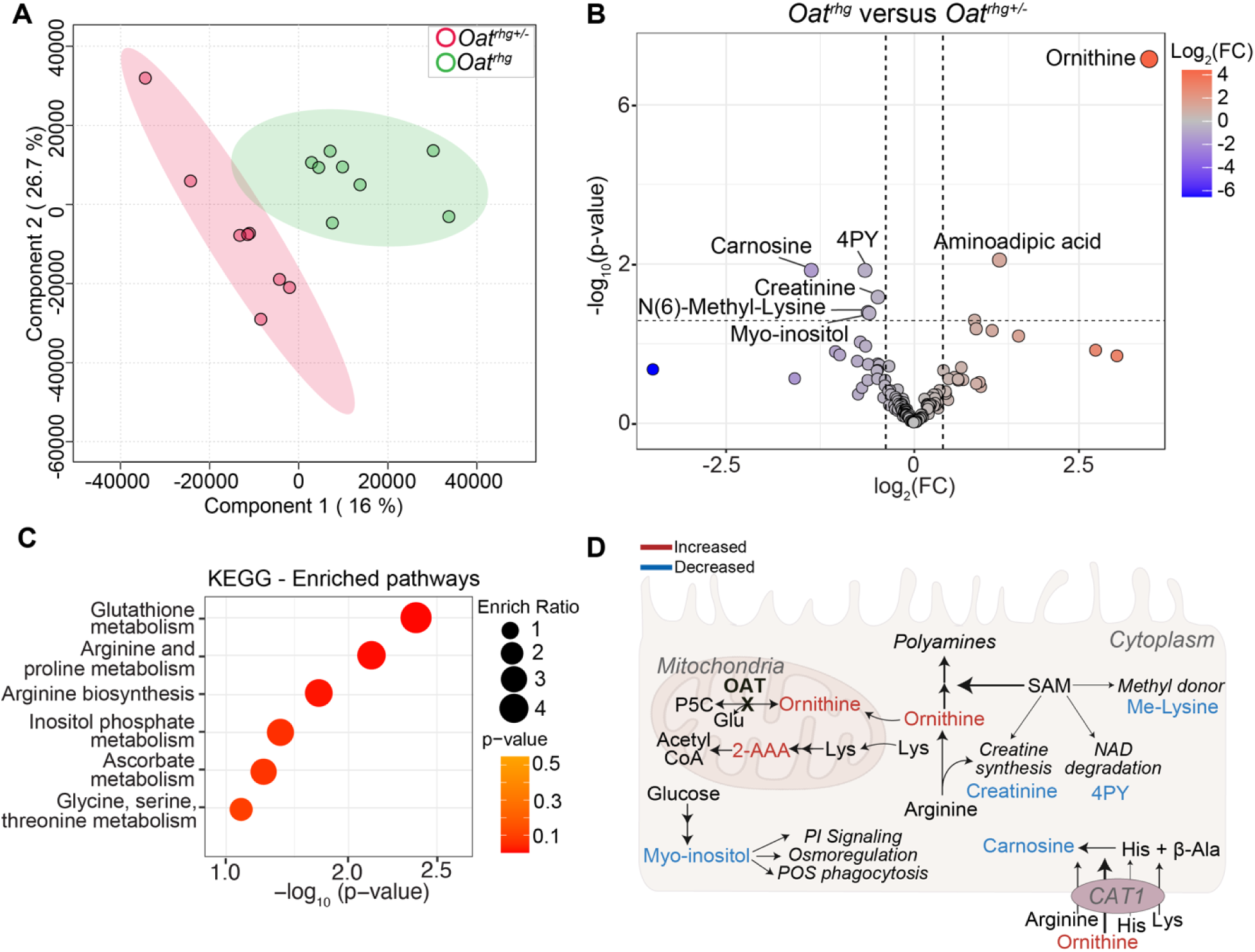
Altered metabolites in RPE/Cho of *Oat^rhg^* compared to controls. **(A)** Partial least squares discriminant analysis (PLS-DA) of RPE/Cho metabolomics distinguishes *Oat^rhg^* from *Oat^rhg+/-^* mice. **(B)** Volcano plot of RPE/Cho metabolomics showing increased (right) and decreased (left) in *Oat^rhg^* relative to *Oat^rhg+/-^*. **(C)** KEGG pathway enrichment analysis of RPE/Cho metabolomics comparing *Oat^rhg^* and *Oat^rhg+/^*^-^. **(D)** Proposed model of metabolomic alterations in *Oat^rhg^* RPE/Cho. Excess ornithine may be redirected toward polyamine synthesis, impacting the SAM pool and downstream pathways. In addition, high ornithine may compete with other cationic amino acids for shared transporters, altering their cellular uptake. Red indicates increased and blue decreased metabolites in *Oat^rhg^* relative to controls. Darker arrows indicate putative enhanced metabolic flux or transport. Pyrroline-5-carboxylate (P5C); ornithine aminotransferase (OAT); Glutamate (Glu); Lysine (Lys); Aminoadipic acid (2-AAA); Coenzyme A (CoA); S-adenosyl methionine (SAM); Me-Lysine (N6-Methyl-Lysine); N1-Methyl-4-pyridone-3-carboxamide (4PY); Histidine (His); β-Alanine (β-Ala); Cationic amino acid transporter 1 (CAT1); Phosphoinositol (PI); Photoreceptor outer segment (POS). N=4. Fold change ≥1.3, p-value < 0.05.

### 3.7. OAT deficiency drives tissue-specific proteomic remodeling with few shared changes across tissues

To investigate common effects of OAT deficiency in tissue proteomes, we compared DE proteins across RPE/Cho, retina, and liver (**Figure 7A**). OAT was the only protein consistently reduced in all three tissues at both ages, confirming the robustness of our model. RPE/Cho showed the most extensive proteomic remodeling, followed by liver and then retina, supporting RPE/Cho as the initial site of damage in OAT deficiency. Aside from OAT, six proteins were consistently altered across all three tissues. Proteins such as TNNT3, CP341, BGAL, and Q8HWB2 changed in the same direction in each tissue, whereas MYH7B and CPLX4 showed opposite regulation between RPE/Cho and liver (**Figure 7B**). These shared proteins are involved in cytoskeletal structure and vesicle trafficking (TNNT3, MYH7B, CPLX4), lysosomal processing (BGAL), immune signaling (Q8HWB2), and lipid metabolism (CP341), highlighting a common cellular response to OAT loss.

**Figure 7.**
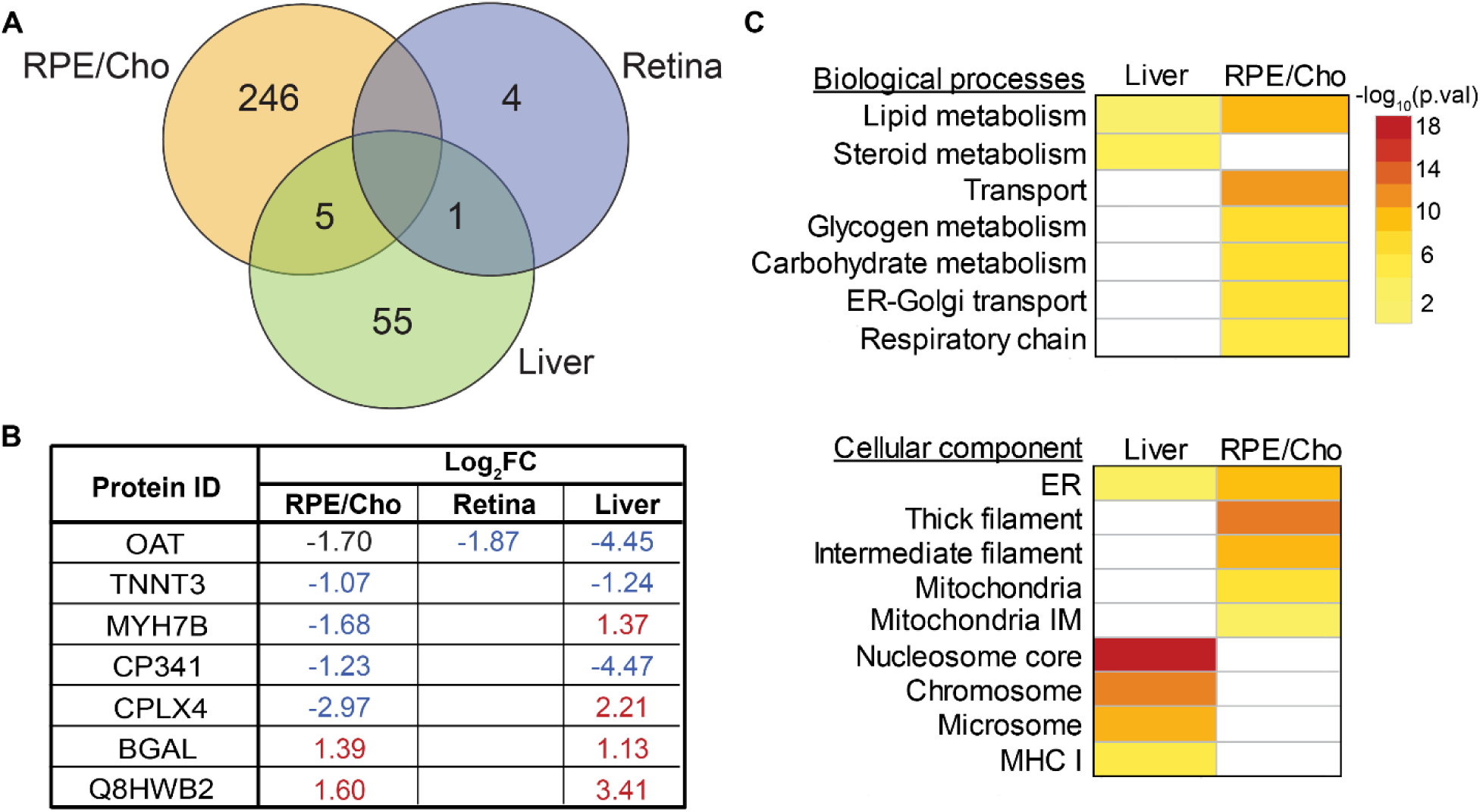
Cross-tissue proteomic overview of DE proteins in *Oat^rhg^* compared to control mice. **(A)** Total significantly changed and overlapping proteins in *Oat^rhg^* in the RPE/Cho, retina, and liver tissues. **(B)** Overlapping DE proteins with the corresponding Log_2_FC, in blue for downregulated, red for upregulated, black for non-significantly changed in *Oat^rhg^* tissues. **(C)** Tissue-specific and overlapping enriched pathways in RPE/Cho and liver proteomics, categorized by biological processes and cellular components. Retina was not included as the number of significantly altered proteins was insufficient for enrichment analysis. Endoplasmic reticulum (ER); Mitochondria inner membrane (IM); Major Histocompatibility Complex I (MHC I). |log_2_fold change| ≥ 1, p-value < 0.05.

We next compared GO-enriched biological processes and cellular components between RPE/Cho and liver (**Figure 7C**). Retina was not included as the number of significantly altered proteins was not sufficient for GO analysis. At the level of biological processes, both tissues showed enrichment of pathways related to lipid metabolism. In RPE/Cho, additional enriched processes were related to transport, carbohydrate metabolism, and the respiratory chain, highlighting broader alterations in RPE/Cho from OAT deficiency. In the cellular component category, both tissues showed overlap in ER-associated proteins. RPE/Cho further showed enrichment of cytoskeletal and mitochondrial components, whereas in the liver, nucleosome-and chromosome-associated proteins were dominant. These differences likely reflect inherent tissue programs, with the liver retaining regenerative capacity, while the post-mitotic RPE/Cho has limited ability to adapt. Overall, these findings indicate that despite minimal overlap in individual proteins and pathways, OAT deficiency drives distinct tissue-specific proteomic changes.

### 3.8. Cross-tissue metabolomic analysis reveals early shared metabolic alterations in OAT deficiency

To identify common OAT⍰dependent metabolite changes across liver, RPE/Cho, and retina, we compared significantly altered metabolites in the three tissues using a Venn diagram (**Figure 8A**). Only two metabolites were shared among all tissues, and four were shared between RPE/Cho and retina (**Figure 8A, B**). Ornithine was consistently increased, whereas N(6)⍰methyl⍰lysine decreased in all tissues, linking ornithine accumulation to altered lysine and methylation pathways. This was further supported by reduced levels of 4PY, a nicotinamide⍰derived metabolite in both RPE/Cho and retina, as well as shared alterations in aminoadipic acid, which was increased in RPE/Cho but decreased in retina. Retina displayed the largest number of altered metabolites (13), compared to 7 in RPE/Cho and liver, suggesting greater metabolic vulnerability to OAT deficiency. KEGG pathway analysis revealed a common disruption of amino⍰acid metabolism, including arginine biosynthesis, arginine and proline metabolism, and glutathione metabolism, in all three tissues (**Figure 8C**). Together, these findings indicate that OAT loss, in addition to altering ornithine⍰related pathways, leads to early disruptions in one-carbon and redox pathways, with retinal metabolism being the most affected.

**Figure 8.**
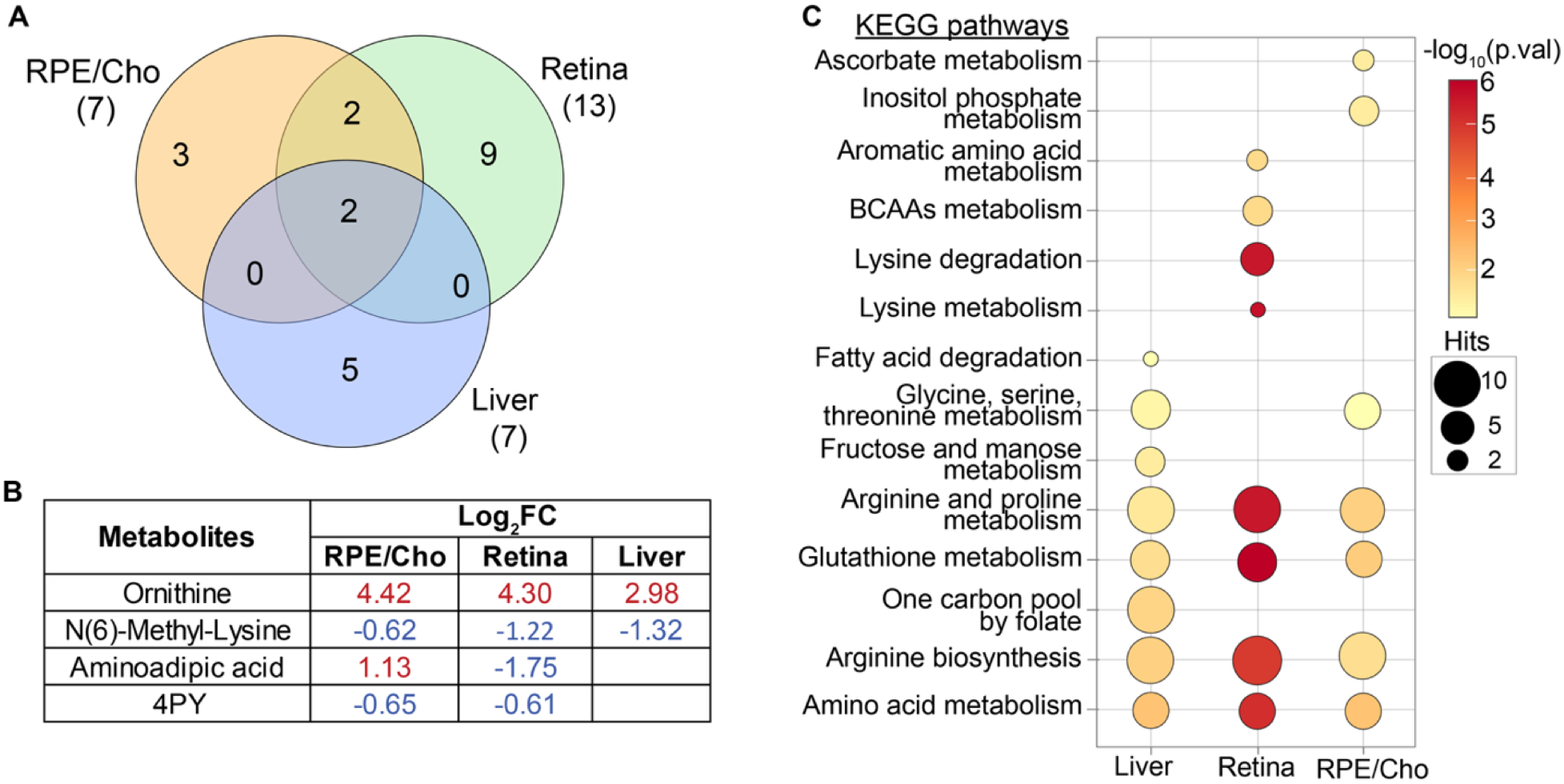
Cross-tissue metabolomic overview of DE proteins in *Oat^rhg^* compared to controls. **(A)** Unique and overlapping significantly changed metabolites across tissues in *Oat^rhg^*and **(B)** showing the overlapping metabolites and their Log_2_FC, in blue for downregulated and red for upregulated. **(C)** Comparison of top enriched metabolic pathways from KEGG analysis in liver, retina, and RPE/Cho. N1-Methyl-4-pyridone-3-carboxamide (4PY). Fold change ≥ 1.3, p-value < 0.05.

## 4. Discussion

In this study, we have identified early metabolic and proteomic signatures of liver and ocular tissues in OAT deficiency. Ornithine accumulation and reduced N(6)-methyl-lysine are common metabolic features across tissues. The liver metabolome reflects increased ornithine disposal through the urea cycle and impaired SAM-dependent pathways, accompanied by proteomic changes in detoxification enzymes and histone proteins. The retinal metabolome highlights reprogrammed amino acid metabolism to support glutamate homeostasis, with few proteomic changes in cytoskeletal and synaptic proteins. In contrast, RPE/Cho shows the most extensive proteomic remodeling, including downregulation of mitochondrial proteins and alterations in cytoskeletal and ECM-associated proteins. These alterations are accompanied by reduced metabolites linked to mitochondrial function and antioxidant capacity, including creatine, lysine, carnosine, and myo-inositol, indicating early and selective vulnerability of the RPE/Cho.

Beyond ornithine accumulation, a common feature of OAT deficiency in both liver and ocular tissues is the disruption of SAM-dependent metabolism, as indicated by reduced N(6)-methyl⍰lysine in all tissues. Excess ornithine can be redirected into polyamine synthesis, an important SAM-consuming pathway that uses decarboxylated S-adenosylmethionine (dcSAM) and may limit SAM availability for methylation reactions [25,70]. While SAM-related metabolites and proteins were consistently affected, their downstream effects were tissue-specific. In the liver, our data suggest metabolic compensation by increasing betaine to support SAM regeneration and upregulation of the histone demethylase KDM3B [71,72]. These changes in histone methylation patterns likely alter chromatin structure, leading to transcriptional reprogramming, including increased expression of histone H2B proteins [73,74]. Moreover, the reduced transsulfuration flux from SAM may impact glutathione levels and sulfate⍰dependent detoxification [71].

In contrast to the liver, the eye metabolome and proteome showed limited evidence of metabolic compensation. Reduced levels of 4PY, a methylated nicotinamide byproduct, in both retina and RPE/Cho indicate decreased SAM-mediated methylation. In addition, retinal cystathionine and xanthosine were diminished further reflecting impaired glutathione synthesis and purine metabolism, both essential for photoreceptor function and survival [75–78]. SAM homeostasis depends on *de novo* synthesis from methionine and recycling from S-adenosyl-L-homocysteine (SAH). In the RPE/Cho, simultaneous downregulation of MAT2B (involved in SAM synthesis) and SAHH2 (SAH hydrolase) suggests a disruption of the methionine cycle. This deficit is corroborated by reduced levels of the SAM-dependent metabolites, including creatinine, 4PY, and N(6)-methyl-lysine. These findings highlight that ocular tissues have limited capacity to activate alternative pathways to compensate for these disruptions.

The liver detoxifies ammonia through the urea cycle by converting ornithine to citrulline, which is then regenerated from arginine to sustain the cycle. Spatially, urea cycle enzymes are enriched in periportal hepatocytes, whereas OAT, together with cytochrome P450 enzymes involved in detoxification, are restricted to perivenous hepatocytes [79–81]. Our proteomic data reveal changes in detoxification proteins that map predominantly to this perivenous zone. The accumulated ornithine and citrulline in the *Oat^rhg^* liver (**Figure 3B**) suggest that periportal activity remains intact, possibly shunting excess ornithine into the urea cycle. This increased urea cycle flux may impose a high bioenergetic demand on the liver to sustain the ATP requirements, consistent with decreases in lactate and fructose. Although the liver supplies key metabolites necessary for retinal health (e.g., vitamin A, taurine, and unsaturated fatty acids) [82–84], it remains to be determined whether such localized disruptions might compromise the availability of factors essential for photoreceptor and RPE survival in GA.

The retinal metabolome reflects an activation of alternative pathways to preserve glutamate. Glutamate functions as the principal excitatory neurotransmitter for phototransduction and a central metabolic intermediate, supporting amino acid metabolism, redox balance, and mitochondrial function [56,85,86]. To maintain glutamate pools, retina has a high cytosolic reducing power to protect glutamate oxidation through the malate-aspartate shuttle [55].

Glutamate can also be regenerated through transamination of α-KG with other amino acids such as alanine, BCAAs, and lysine. Strikingly, BCAAs, lysine, and its derivative, aminoadipic acid, are reduced in *Oat^rhg^* retinas. Catabolism of these amino acids produces glutamate and yields succinyl-CoA (from valine) and acetyl-CoA (from leucine and lysine) to support mitochondrial metabolism (**Figure 4C, E**). Importantly, entry through succinyl-CoA in TCA bypasses α-KG, which can help preserve the α-KG/glutamate pool required for the malate-aspartate shuttle in retinal neurons.

RPE/Cho shows the most substantial proteomic dysregulation, specifically in mitochondrial metabolism, cytoskeletal, and ECM-related proteins (**Figure 5F**). These altered protein expressions may compromise cell bioenergetics, polarity, transport, and barrier integrity, processes closely linked to RPE dysfunction in retinal disease [87–92]. The mitochondrial proteins involved in oxidative phosphorylation (e.g., NDUBB, NDUS8, COX2), transport (TOM22, MPC1), and fusion (MFN1) are all downregulated, suggesting early mitochondrial dysfunction in *Oat^rhg^* RPE/Cho. Additionally, reduced creatinine and downregulation of mitochondrial creatine kinase (KCRS) further support mitochondrial dysfunction in RPE. Creatinine is a degradation product of creatine, a crucial molecule to support stable ATP supply in energetically demanding tissues via creatine kinase [93,94]. Elevated ornithine is known to inhibit AGAT, the rate-limiting enzyme for creatine synthesis, which may explain reduced creatine availability [20]. These findings align with mitochondrial abnormalities reported in GA patients [95–97].

RPE requires robust mitochondria to oxidize nutrients derived from the daily phagocytosis of photoreceptor outer segments and to produce nutrients that support the neural retina [98]. Mitochondrial dysfunction in RPE is sufficient to induce RPE dedifferentiation, ECM remodeling, and photoreceptor degeneration [99–101]. Unlike other cells, RPE preferentially utilizes proline as a fuel to support mitochondrial metabolism and amino acid synthesis for both RPE and the neural retina [57,98]. We speculate that loss of OAT in RPE/Cho may limit proline synthesis by a dual mechanism: reduced proline-derived ornithine through P5C and inhibition of pyrroline-5-carboxylate synthase (P5CS) due to accumulated ornithine [102]. However, proline levels remained stable in the *Oat^rhg^* RPE/Cho. This is likely compensated by reduced proline oxidation in part due to impaired mitochondria, while simultaneously decreasing proline utilization for ECM and collagen synthesis. Consistently, several ECM proteins are altered in RPE/Cho proteomics. Interestingly, proline supplementation has been reported to slow or halt the progression of chorioretinal lesions in GA [29]. Further studies are needed to assess the role of proline in RPE mitochondrial dysfunction and protection in GA.

In addition, the decreased levels of carnosine and myo-inositol further support impaired RPE metabolism. Carnosine (β-alanyl-L-histidine) is a mitochondria-protective dipeptide that scavenges reactive oxygen species and limits lipid peroxidation [68,103,104]. Its decline suggests reduced antioxidant capacity and compromised barrier integrity in the RPE, further exacerbating mitochondrial function [69,105,106] (**Figure 6B, D**). Myo-inositol is a precursor for phosphatidylinositol (PI) synthesis. Strikingly, CDP-diacylglycerol-inositol 3- phosphatidyltransferase (CDIPT), a key enzyme for the synthesis of PI, was downregulated in *Oat^rhg^* RPE/Cho (**Table S10**). Since phosphoinositides are important for regulating membrane trafficking, cytoskeletal organization, mitochondrial function, and outer segment phagocytosis [65–67,107–110], their downregulation may contribute to the cytoskeletal and mitochondrial defects in *Oat^rhg^* RPE/Cho.

Together, our data demonstrate that OAT deficiency drives early, common, and tissue-specific changes in the metabolome and proteome that extend beyond ornithine metabolism. These findings support the need for therapeutic strategies in GA that target both systemic metabolism and tissue-specific pathways. Approaches aimed at lowering plasma ornithine, together with ocular tissue-targeted metabolic or gene therapies, may be required to preserve visual function and potentially benefit other affected tissues in GA, including the cornea, lens, muscle, and brain [7,9,15,20,111,112].

While our study provides an early molecular signature of liver and eye in OAT deficiency, it has some limitations. First, proteins were sampled from a single hepatic region, which may not capture the full cellular and functional tissue heterogeneity, particularly given the spatial organization of urea cycle enzymes and OAT [80,113,114]. Second, the RPE and choroid were collected together to better preserve RPE integrity. Future studies are required to dissect their tissue-specific changes by optimized separation techniques, staining, or spatial proteomics.

Third, although OAT expression and activity were experimentally validated, the mechanistic roles of other altered proteins and their associated biochemical pathways are currently inferred and await direct functional validation.

## Author Contributions

Conceptualization: Artjola Puja, Jianhai Du

Methodology: Artjola Puja, Isabella Mascari, Tuan Ngo, Rong Xu

Investigation (experiments/data collection): Artjola Puja, Rong Xu, Isabella Mascari, Meghashri Saravanan

Resources (animals/samples/instrumentation): Artjola Puja, Isabella Mascari, Tuan Ngo, Rong Xu, Qingyan Wang, Ying Zhang

Visualization (figures/tables): Artjola Puja, Meghashri Saravanan

Writing – Review & Editing: Artjola Puja, Isabella Mascari, Tuan Ngo, Rong Xu, Qingyan Wang, Ying Zhang

Supervision: Jianhai Du

## Funding Acquisition

NIH Grants (EY026030, EY031324, EY032462), the Retina Research Foundation, NIH/NIGMS grant R24GM137786 to IDeA National Resource for Quantitative Proteomics, NIH NIGMS P20GM144230 Visual Sciences COBRE grant to WVU, and an unrestricted challenge grant from Research to Prevent Blindness (RPB) to the Ophthalmology department at WVU.

## Acknowledgements

This work was supported by NIH Grants (EY026030, EY031324, EY032462), the Retina Research Foundation, NIH/NIGMS grant R24GM137786 to IDeA National Resource for Quantitative Proteomics, NIH NIGMS P20GM144230 Visual Sciences COBRE grant to WVU, and an unrestricted challenge grant from Research to Prevent Blindness (RPB) to the Ophthalmology department at WVU.

We also thank Dr. David Sean Hansman, postdoctoral fellow in our laboratory, for reviewing the manuscript and providing valuable suggestions.

## Data availability

The mass spectrometry proteomics raw data have been deposited to the ProteomeXchange Consortium via PRIDE partner repository with the dataset identifier PXD063614 and 10.6019/PXD063614. The metabolomics raw data have been deposited to MassIVE repository with identifier MSV000101103.

## Abbreviations

GA: gyrate atrophy of the choroid and retina
OAT: ornithine aminotransferase
P5C: pyrroline-5-carboxylate
PLP: Pyridoxal 5’-posphate
RPE/Cho: retinal pigment epithelium/choroid
TCA cycle: tricarboxylic acid cycle
ER: endoplasmic reticulum
ECM: extracellular matrix
GO: Gene Ontology
DE: differentially expressed
BCAA: branched chain amino acid
SAM: S-adenosyl methionine
SAH: S-adenosyl homocysteine.

