## Supplementary Figures for "Early Proteomic and Metabolic Signatures of Liver and Eye in OAT-Deficient Mice"

This PDF file includes:

Supplementary Figures

Supplementary Methods

Key Resources Table

**Supplementary Figures**

**Supplementary Figure 1**

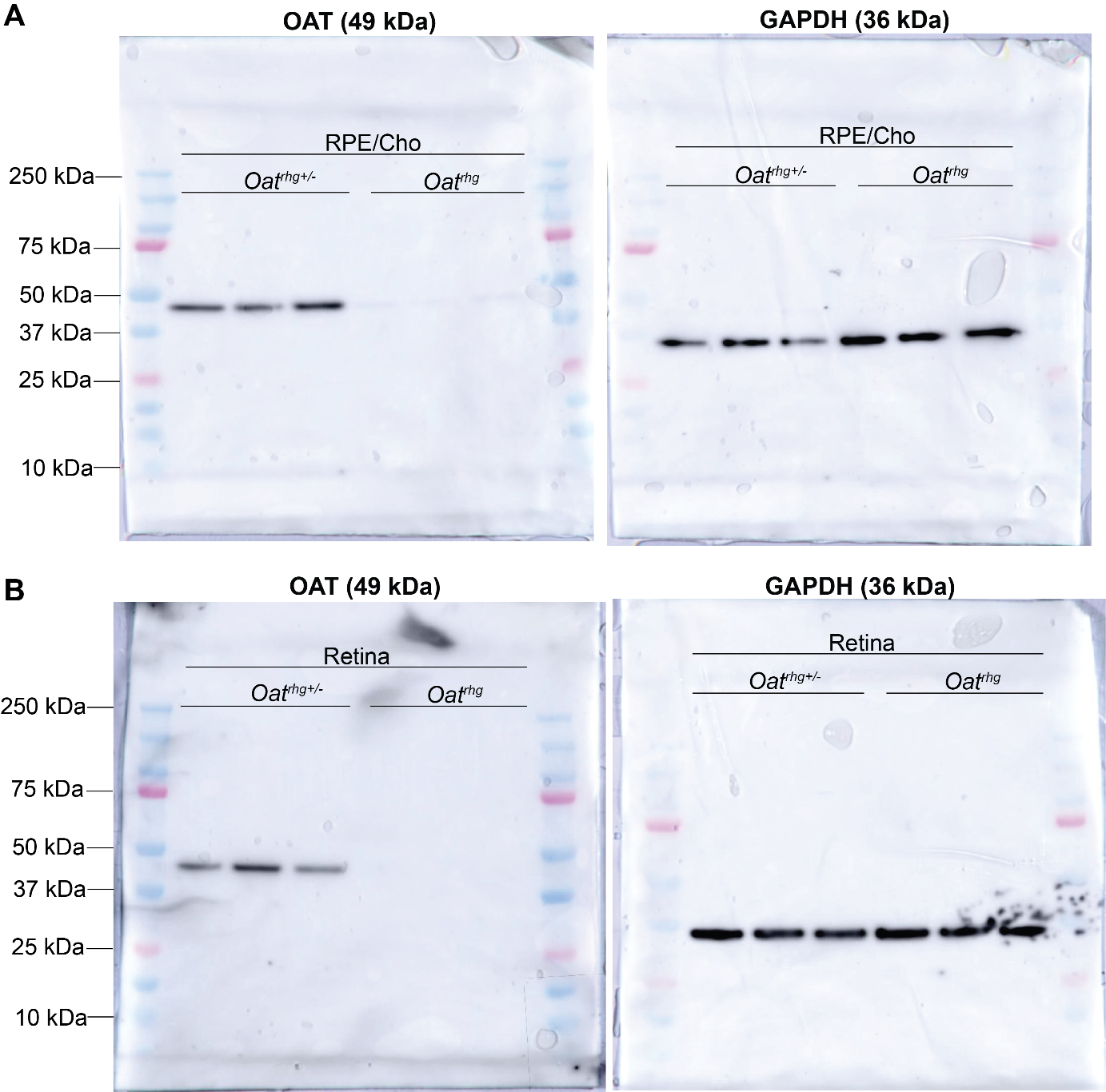

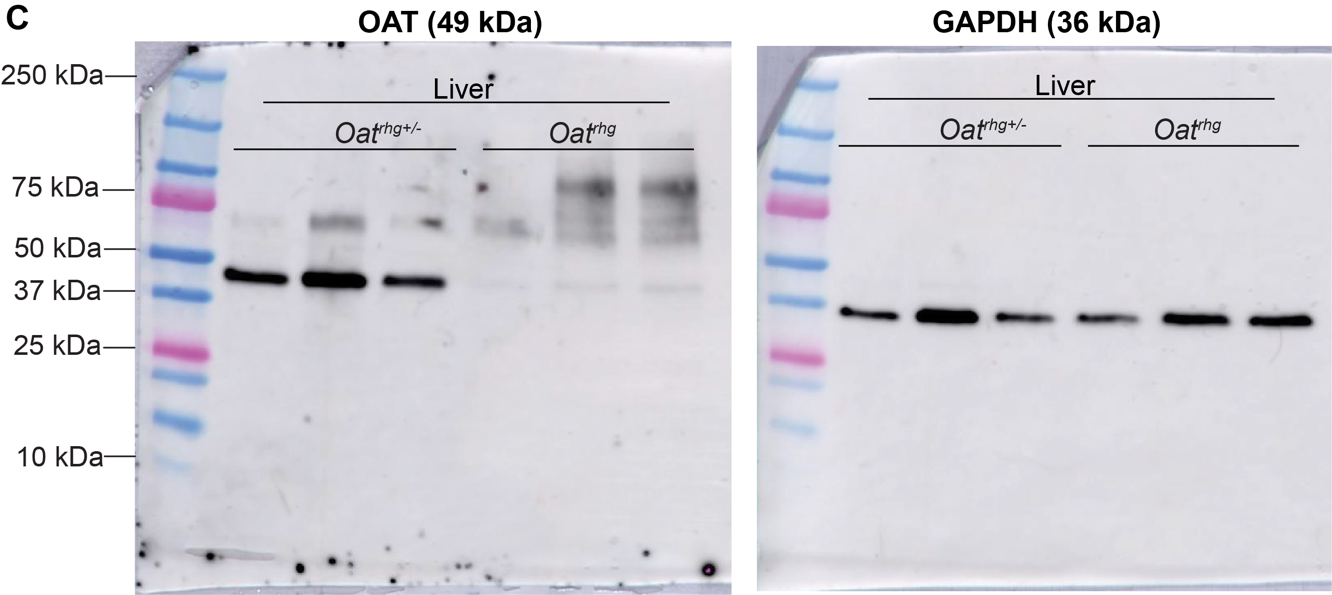

**Figure S1. Raw data for Western blots included in the Figure 1C.** (**A-C**) OAT protein expression in RPE/Cho, retina, and liver, respectively from heterozygous controls *Oat^rhg+/-^* and null mutant *Oat^rhg^* mice. GAPDH was used as loading control.

**Supplementary Figure 2**

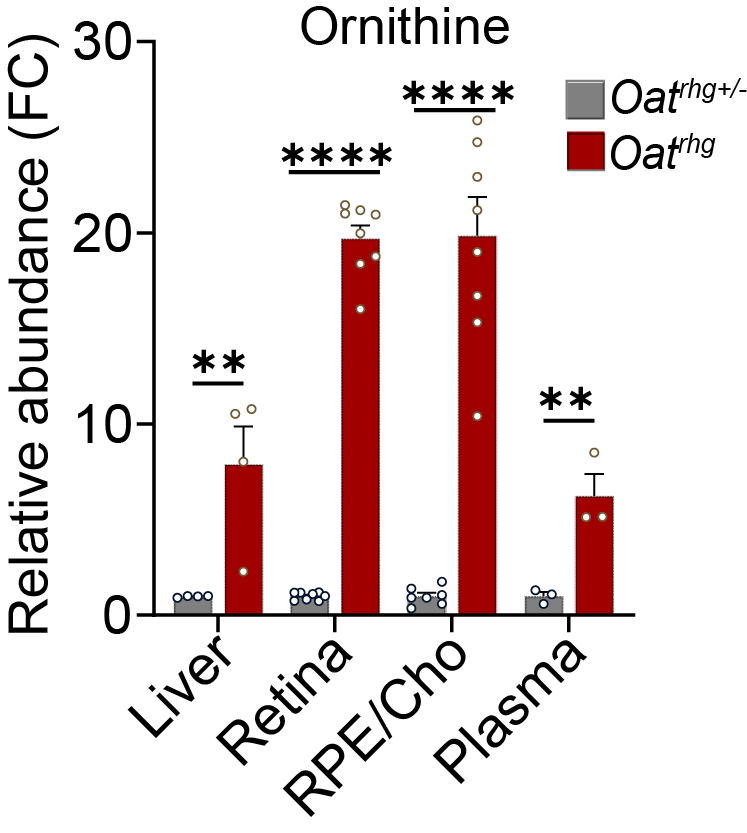

**Figure S2. OAT deficiency leads to systemic accumulation of ornithine across tissues and plasma.** Bar graphs show ratios of ornithine abundance in liver, retina, RPE/Cho, and plasma in *Oat^rhg^* relative to *Oat^rhg+/-^*. p-value < 0.05. RPE/Choroid (RPE/Cho).

**Supplementary Figure 3**

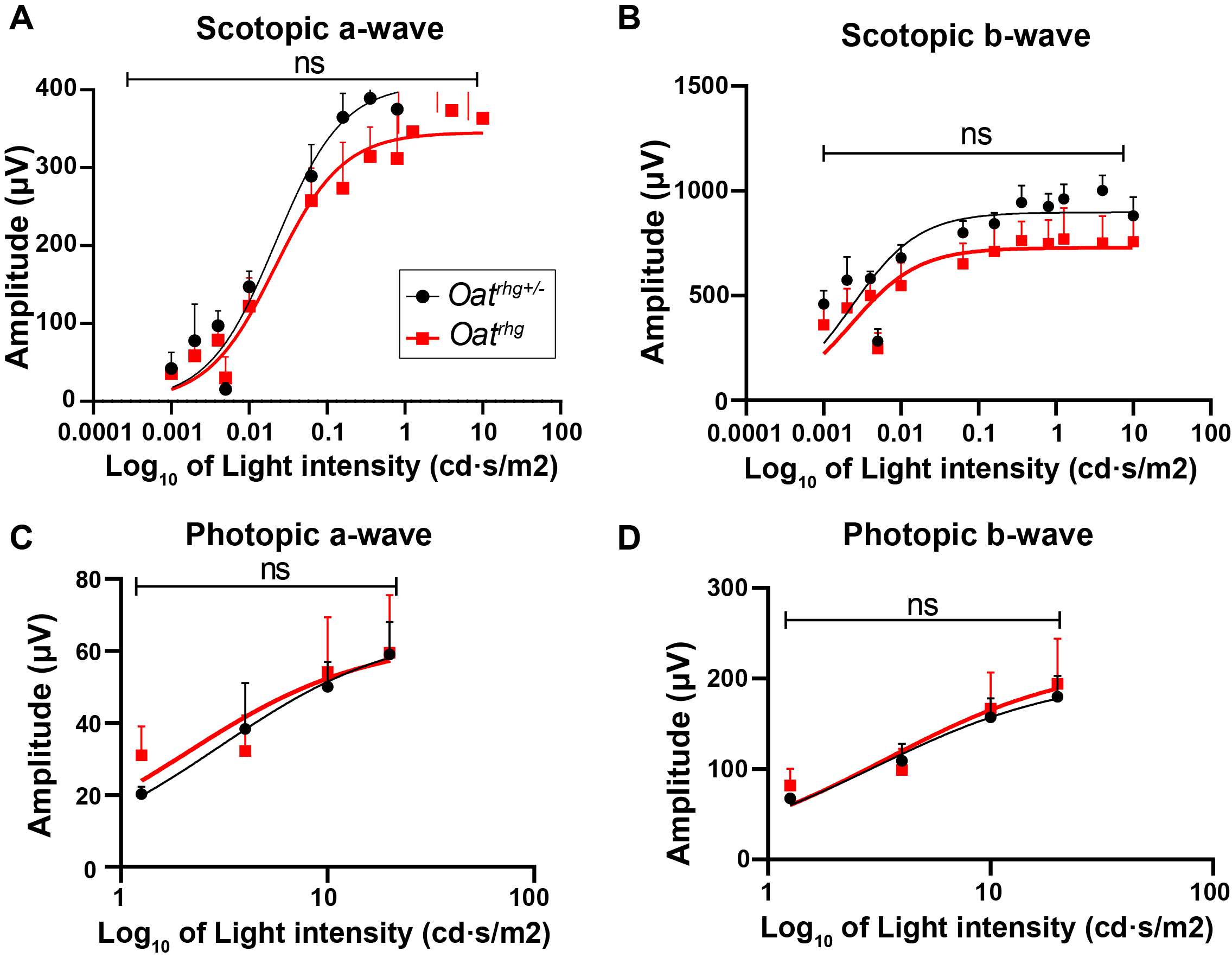

**Figure S3. Analysis of the visual function by electroretinogram (ERG) in *Oat^rhg+/-^* and *Oat^rhg^*.** ERG responses of increasing intensity were obtained for *Oat^rhg+/-^* and *Oat^rhg^* mice at around 3 months of age, under dark-adapted (scotiopic; **A-B**) and light-adapted (photopic; **C-D**) eyes. Intensity of mouse response is plotted as a function of amplitude for a- and b-waves versus varying log10 of light intensities. t-test analysis using GraphPad v.9.5.1, did not reveal any statistically significant difference between *Oat^rhg^* compared to *Oat^rhg+/-^*. Non-significant (ns); p-value < 0.05, N=3.

**Supplementary Figure 4**
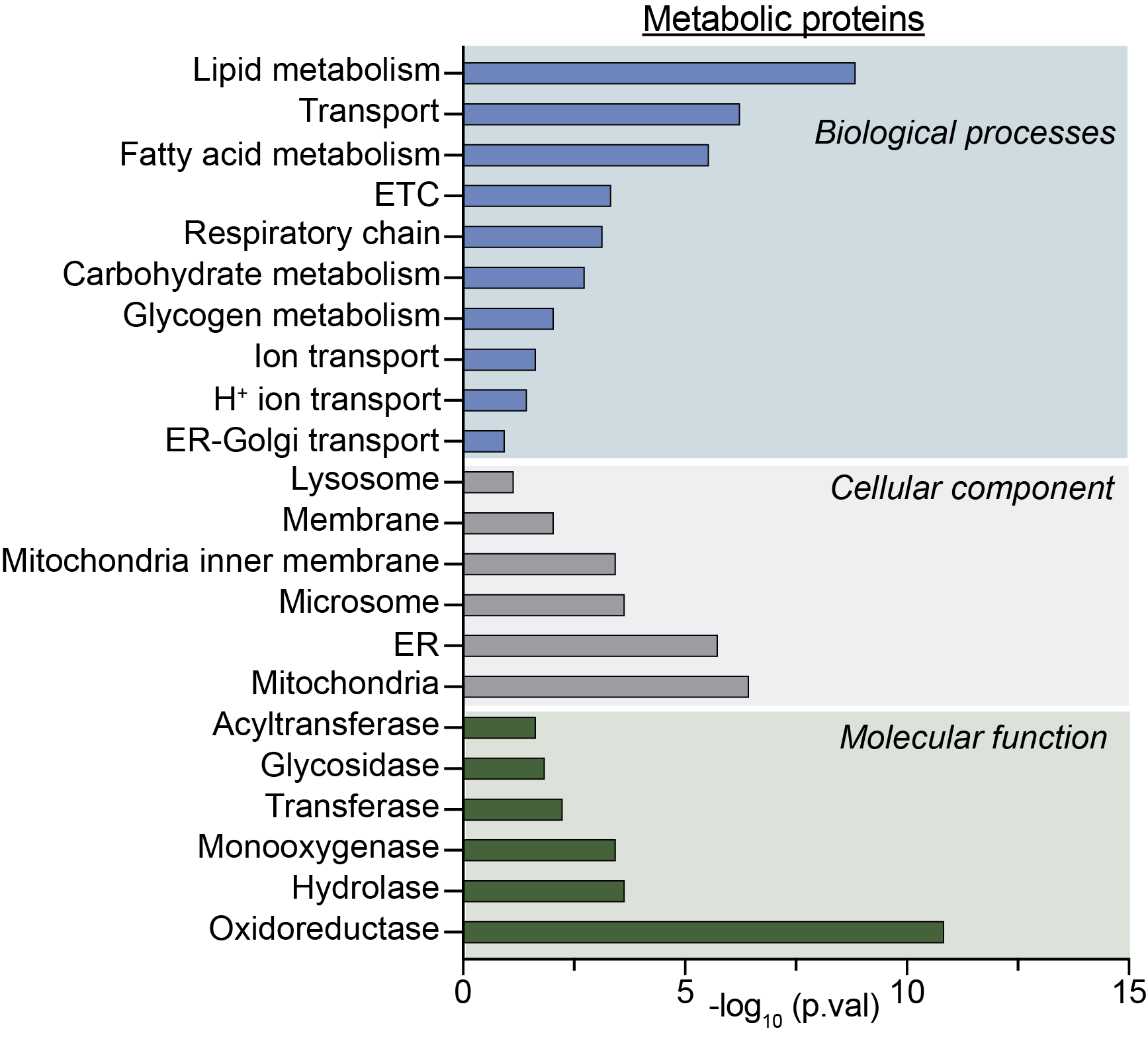
 **Figure S4. Functional annotation analysis of enriched metabolic proteins in RPE/Cho *Oat^rhg^*.** Functional enrichment analysis was performed using DAVID and functional annotation categories including biological processes, cellular components, and molecular functions are shown. Significance was determined by |log_2_fold change| ≥ 1, p-value < 0.05. Electron transport chain (ETC); Endoplasmic reticulum (ER).

**Supplementary Methods

Electroretinogram (ERG)**

*Oat^rhg+/-^* and *Oat^rhg^* mice were dark-adapted overnight prior to electroretinography (ERG). ERG recordings were obtained using the Celeris System (Diagnosys LLC, Lowell, MA) equipped with a two-channel amplifier, dual function stimulator-light guide electrode and Espion software (version 6; Diagnosys). Mice were anesthetized in an induction chamber with 5% isoflurane in 2.5 L/min oxygen for one minute, then transferred to a heated platform set at 37 °C. Anesthesia was maintained with 1.5% isoflurane in 2.5 L/min oxygen delivered via nose cone. Pupils were dilated for 10 minutes using Tropi-Phen (a 1:1 mixture of 2.5% phenylephrine and 1% tropicamide) from Pine Pharmaceuticals. Two platinum subdermal electrodes were placed, one serving as the reference electrode in the temporal scalp region between the eye and ear (inserted 1–2 mm deep at a shallow angle to remain subdermal), and the other as the ground electrode in the hind limb (inserted 1–2 mm deep). Two corneal electrodes are positioned on the surface lubricated with Systane Ultra Lubricant Eye Drops (Alcon). All procedures were conducted under the dime red light. Scotopic ERG responses were recorded at flash intensities of 0.0005–10 cd·s·m⁻², with responses averaged at each intensity. Mice were then light-adapted for 10 minutes, after which photopic responses were recorded using flashes of 1.26–20 cd·s·m⁻². Data were normalized to baseline, and each eye was analyzed separately to quantify A-wave and B-wave amplitudes, which were then averaged and graphed.

**Table S1. Key resources**

| **Mouse Strain** | **Catalog** | **Company** | **Location** |
| --- | --- | --- | --- |
| B6Ei; AKR-*Oat^rhg^*/J | RRID: IMSR_JAX:003544 | The Jackson Laboratory | Bar Harbor, ME, USA |
| **Primer sequence** | **Catalog** | **Company** | **Location** |
| F: 5’ AGGCCCTGAAAATG AAAAGC 3’ |  | Integrated DNA Technologies | Coralville, IA, USA |
| R: 5’ TTTGAAAGGGCTCTC CACAC 3’ |  | Integrated DNA Technologies | Coralville, IA, USA |
| **Reagents** | **Catalog** | **Company** | **Location** |
| RIPA Lysis and Extraction Buffer | 89901 | Thermo Fisher Scientific | Rockford, IL USA |
| Pierce Protease and Phosphatase Inhibitor Mini Tablets | A32959 | Thermo Fisher Scientific | Rockford, IL USA |
| Pierce™ BCA Protein Assay Kit | 23225 | Thermo Fisher Scientific | Rockford, IL USA |
| Tween-20 | 9005-64-5 | Millipore Sigma | Burlington, MA USA |
| 4–20% Mini-PROTEAN^®^ TGX™ Precast Protein Gels | 4561094 | Bio-Rad | Hercules, CA USA |
| 10x Tris Buffered Saline (TBS) | 1706435 | Bio-Rad | Hercules, CA USA |
| 10x Tris/Glycine/SDS Buffer | 1610772 | Bio-Rad | Hercules, CA USA |
| Blotting-Grade Blocker (non-fat dry milk) | 1706404 | Bio-Rad | Hercules, CA USA |
| Immobilon Western Chemiluminescent HRP Substrate | WBKLS0500 | Millipore Sigma | Merck KGaA, Darmstadt, Germany |
| Phosphate buffer saline | P3813-10PAK | Sigma-Aldrich | St. Louis, MO USA |
| Methanol, Optima™ LC/MS Grade | A456-4 | Fisher Scientific | Waltham, MA USA |
| Methoxyamine hydrochloride | 226904-1G | Sigma-Aldrich | St. Louis, MO USA |
| N-tert-butyldimethylsilyl-N-methyltrifluoroacetamide (TBDMS) | 190500 | Sigma-Aldrich | St. Louis, MO USA |
| Systane Lubricant Eye Gel |  | Alcon | Fort Worth, TX USA |
| Tropi-Phen (Tropicamide 1%: Phenylephrine HCL 2.5%) | 69194-762-01 | Pine Pharmaceutical | Tonawanda, NY USA |
| Water Optima^TM^ LC/MS Grade | 166415 | Fisher Scientific | Waltham, MA USA |
| Methanol | 164905 | Fisher Scientific | Waltham, MA USA |
| Acetonitrile, Optima^TM^ LC/MS Grade | 75-05-8 | Fisher Scientific | Waltham, MA USA |
| Ammonium Acetate | 431311 | Sigma-Aldrich | St. Louis, MO USA |
| **Antibodies & Host** | **Catalog** | **Company** | **Dilutions** |
| Anti-OAT Antibody, mouse | sc-374243 | Santa Cruz | 1:1000 (Western Blot) |
| Anti-GAPDH Antibody, rabbit | 5174S | Cell Signaling | 1:1000 (Western Blot) |
| Rat anti-Mouse IgG HRP | ab131368 | Abcam | 1:2000 (Western Blot) |
| Anti-Rabbit IgG HRP | 7074S | Cell Signaling | 1:2000 (Western Blot) |
